# Plant-specific *tau* class glutathione transferases safeguard against lipid oxidation stress through dual detoxification of lipid peroxides and reactive carbonyl species

**DOI:** 10.1101/2025.11.21.687491

**Authors:** Yasuo Yamauchi, Nagisa Matsuura, Fumiya Kawaguchi, Chisato Sakurai, Ryosuke Nakasuga, Junya Ito, Kiyotaka Nakagawa, Masaharu Mizutani, Yukihiro Sugimoto, Jun’ichi Mano

## Abstract

- Oxidation of membrane lipids results in the formation of lipid peroxides (LOOHs) and reactive carbonyl species (RCS), that induce lipid oxidation stress in plants. The plant-specific *tau* class of glutathione transferases (GSTUs) possess both LOOH peroxidase and RCS-conjugating activities, but their overall contribution to stress tolerance remains unclear.
- We systematically examined 16 GSTU isozymes of *Arabidopsis thaliana* and found that all catalysed glutathione-dependent reduction of LOOHs. GSTU17, showing the highest activity, recognized five different isomers of linoleic acid-derived hydroperoxides within comparable catalytic efficiencies.
- T-DNA mutants lacking GSTU3, GSTU17, GSTU18, GSTU19 or GSTU25—isoforms with strong conjugation activities toward acrolein, 4-hydroxynonenal or crotonal—displayed enhanced sensitivity to photooxidative stress, indicating that detoxification of RCS is crucial for oxidative protection.
- Transcriptome analyses revealed that exposure to RCS (crotonal, 2-pentenal or 2-hexenal) strongly induced 12 *GSTU* genes, while other GST classes showed little response. Collectively, these results demonstrate that GSTUs function as a major enzymatic system mitigating lipid oxidation stress in *A. thaliana* through the combined detoxification of lipid peroxides and their reactive carbonyl products.

## Introduction

Formation of reactive oxygen species (ROS) is intrinsically associated with aerobic metabolism. In plant cells, ROS are generated as by-products of energy metabolism in chloroplasts and mitochondria constitutively, and their production increases under environmental stress conditions due to over-reduction of the electron transport chains (Asada 2006). Prolonged exposure of plants to stress conditions causes inactivation of antioxidant enzymes and consumption of antioxidants beyond their regeneration, gradually decreasing cellular antioxidant capacity and elevating steady-state ROS levels, a state defined as oxidative stress (Mano, 2002).

ROS not only directly oxidise proteins and nucleic acids, leading to their denaturation, but also initiate oxidation of membrane lipids to lipid peroxides (LOOHs). Accumulation of these LOOHs leads to disintegration of biological membranes and modify their fluidity and permeability (Wong-ekkabut et al. 2007), thereby imposing stress on the cell (Gaschler & Stockwell, 2017). LOOHs can further decompose to yield secondary reactive molecules known as reactive carbonyl species (RCS) or electrophilic oxylipin derivatives (Mano, 2012; Farmer & Mueller, 2013) are generated. RCS are a collective term for aldehydes and ketones with *α*,*β*-unsaturated bonds, including representative species such as acrolein, 4-hydroxynonenal (HNE), and malondialdehyde (MDA) (Esterbauer, Schaur & Zollner, 1991). These highly electrophilic molecules form covalent bonds with Lys, His, and Cys residues of proteins and thereby alter – either impair or activate – their functions (Schopfer, Cipollina & Freeman, 2011; Wible & Sutter, 2017; Parvez et al., 2018). In plants, protein modification by RCS contributes significantly to protein carbonylation, alongside direct oxidation of amino acid residues by ROS (Matamoros et al., 2018). Under salt stress, protein modification with RCS increases with the progression of damage (Mano et al., 2014). RCS act as mediators of ROS cytotoxicity in plants (Mano, Biswas & Sugimoto, 2019; Biswas & Mano, 2021). First, transgenic plants overexpressing 2-alkenal reductase (AER; EC 1.3.1.74), an enzyme that specifically reduces the *α*,*β*-unsaturated bonds of RCS (Mano et al., 2002), exhibit enhanced tolerance to high-light stress (Mano et al., 2005), salt stress (Papdi et al., 2008; Sultana et al., 2024), and aluminium stress (Yin et al., 2010). In these overexpressors, the stress-induced accumulation of RCS – particularly acrolein, (*E*)-2-hexenal, and HNE – was suppressed. Second, *Arabidopsis thaliana* mutants deficient in alkenal/one reductase (AOR; Yamauchi et al. 2011), which has similar catalytic activity to AER, showed greater RCS accumulation and photosystem II (PSII) inactivation under photo-oxidative stress than wild-type plants (Yamauchi et al., 2012). Third, exogenous application of carnosine, a dipeptide that detoxifies RCS, suppressed stress-induced RCS accumulation and alleviated damage caused by salt stress (Sultana et al., 2022). Since RCS are primarily generated from LOOHs, the accumulation of LOOHs themselves constitutes an additional stress factor. Thus, the combined impact of LOOHs and RCS can be collectively considered as *lipid oxidation stress*.

To counteract lipid oxidation stress, plants are equipped with multiple mechanisms to scavenge LOOHs and RCS. LOOHs can be enzymatically reduced by peroxidases, including glutathione peroxidases (Herbette et al., 2002), thioredoxin-dependent glutathione peroxidase (Iqbal et al., 2006), plastidic peroxiredoxins (notably 2-Cys Prx and PrxQ) (Lamkemeyer et al., 2006) and a subset of glutathione transferases (GSTs), which catalyze glutathione (GSH)-dependent reduction of organic peroxides (Bartling et al., 1993; Dixon et al., 2009). For detoxifying RCS, plants have several types of oxidoreductases, i.e. AER, AOR, aldo-keto reductases, aldehyde dehydrogenases and aldehyde oxidases (Mano, Biswas & Sugimoto, 2019). In addition, several GST isozymes from *A. thaliana* and spinach scavenge RCS via the GSH conjugation (Dixon et al. 2009; Mano et al., 2017, 2019). Participation of GSTs in the mitigation of lipid oxidation stress *in planta* is also suggested from the observation that the enhanced GSH content lowers the RCS levels in *A. thaliana* (Yin et al. 2017).

In this study, we sought to elucidate how plant GSTs contribute to the mitigation of lipid oxidation stress. GSTs are ubiquitous enzymes that form multigene families across diverse taxa (Sheehan et al., 2001). Among the plant GSTs, the *tau* class (GSTUs) represents the most diversified group and is uniquely restricted to terrestrial plants (Dixon et al., 2002; Monticolo et al., 2017; Micic et al., 2024). The *A. thaliana* genome encodes 28 *GSTU* genes—more than half of all *GST* genes in this species. GSTs are capable of catalysing two distinct detoxification reactions: conjugation of GSH with RCS (RCS-conjugation activity) and GSH-dependent reduction of LOOHs (LOOH peroxidase activity) (Sylvestre-Gonon et al., 2019). Bartling et al. (1993) reported that a GST isozyme of *A. thaliana* shows peroxidase activity toward hydroperoxy-octadecadienoic acid (HPODE) and hydroperoxy-octadecatrienoic acid (HPOTE), hydroperoxides derived from linoleic and linolenic acids, respectively. Gronwald & Plaisance (1998) showed that two isoforms of GST purified from sorghum have peroxidase activity toward HPODE and HPOTE. Dixon et al. (2009) reported that most *tau*, *phi* and *theta* isoforms of *A. thaliana* display peroxidase activity toward cumene hydroperoxide (cumene-OOH; a model organic peroxide) and that theta-class isoforms show particularly high activity toward HPODE and HPOTE. However, despite the numerical predominance and stress inducibility of GSTUs (Sappl et al., 2009; Gullner et al., 2018; Micic et al., 2024), their peroxidase activities toward physiological LOOHs have not been comprehensively investigated. Given their abundance and strong translational activation under oxidative stress, GSTUs are plausible candidates for the enzymatic reduction of LOOHs—the direct precursors of RCS—and may play pivotal roles in stress tolerance through dual detoxification of LOOHs and RCS.

With respect to RCS conjugation, human alpha-, pi-, and mu-class GSTs (A1-1, P1-1, and M1-1, respectively) efficiently catalyse the conjugation of GSH with acrolein (Berhane et al., 1994). Although these classes are absent in plants, comparable conjugation activity toward acrolein is widely detected in angiosperms, including spinach, *A. thaliana*, napa cabbage, rice, onion, pepper, garland chrysanthemum and celery (Mano et al., 2019). We previously demonstrated that one GSTU isoform from spinach (Mano et al. 2017) and at least 10 GSTUs from *A. thaliana* catalyse GSH conjugation with acrolein (Mano et al., 2019). Notably, GSTUs show relatively narrow substrate specificity toward different types of RCS, in contrast to the broad specificity of RCS-detoxifying oxidoreductases (Mano et al. 2019). Despite accumulating biochemical data, the relative contribution of the RCS-conjugating and LOOH peroxidase activities of individual GST isoforms to plant defence against lipid peroxidation stress remains to be elucidated.

In the present study, we systematically examined 16 *A. thaliana* GSTU isoforms to clarify their enzymatic roles in the detoxification of LOOHs and RCS. All recombinant GSTUs exhibited peroxidase activity toward linoleic acid hydroperoxides (HPODEs), and the expression of 12 GSTU genes was induced by treatment with RCS in planta. Mutant plants lacking these GSTU isoforms displayed enhanced sensitivity to oxidative stress compared with the wild-type. Collectively, these results demonstrate that GSTU isoforms have evolved as a major enzymatic system mitigating lipid oxidation stress through the coordinated detoxification of LOOHs and their reactive carbonyl derivatives.

## Materials and Methods

### Chemicals

Acrolein was prepared from acrolein diethylacetal (Tokyo Chemical Industry, Tokyo, Japan) and HNE from HNE dimethylacetal (Sigma-Aldrich Japan, Tokyo, Japan) via acid hydrolysis, followed by neutralization (Mano et al., 2009). Glutathione reductase was obtained from Fujifilm Wako Pure Chemical (Osaka, Japan). All other chemicals were of analytical grade.

### Preparation of lipid peroxides

HPODE isomers were prepared as previously described, with slight modifications (Ito et al., 2015a, b). Linoleic acid (10 g) was dissolved in chloroform (2000 mL) containing 10 μM methylene blue and subjected to photo-oxidation under LED light (approximately 50,000 lux, 48 h, 25 °C). After the reaction, the solution was purified using a silica gel column to obtain a mixture of six HPODE isomers (designated ’HPODE mix’): 9-10*E*,12*E*-HPODE (9EE-HPODE), 9-10*E*,12*Z*-HPODE (9EZ-HPODE), 10-8*E*,12*Z*-HPODE (10-HPODE), 12-9*Z*,13*E*-HPODE (12-HPODE), 13-9*E*,11*E*-HPODE (13EE-HPODE) and 13-9*Z*,11*E*-HPODE (13ZE-HPODE) (see Supporting Information Fig. S1 for structures). Each isomer was further purified using a semipreparative LC-UV system (Shimadzu, Kyoto, Japan) following established protocols (Ito et al., 2015a, b). Product identity was confirmed by mass spectrometry, and mole amounts were determined on a dry-weight basis.

### Plant materials

Seeds of *A. thaliana* GSTU-deficient T-DNA insertion lines (Col-0 background) were obtained from the Arabidopsis Biological Resource Center (Ohio State University, Columbus, OH, USA). Line IDs were SALK_062920C (*gstu2*), SALK_17999 (*gstu3*), SALK_143928C (*gstu4*), SALK_107148 (*gstu5*), SALK_207079C (*gstu6*), SALK_086642C (*gstu7*), SALK_139615C (*gstu17*), SALK_096297C (*gstu18*), SALK_041942 (*gstu19*), SALK_034472C (*gstu24*) and SALK_042213 (*gstu25*). Seeds were sown on moistened Jiffy-7 peat pellets (Sakata Seed Co., Yokohama, Japan). Following 3 d vernalization at 4 °C in the dark, plants were grown at 23 °C under a 14 h light/10 h dark photoperiod (80 µmol photons m^−2^ s^−1^) without additional nutrients. Homozygous lines were confirmed by genomic DNA PCR with gene-specific primers (Supporting Information Table S1). For oxidative stress treatments, 3-week-old plants were exposed to high light (1,800 µmol m^−2^ s^−1^) with or without 100 µM methyl viologen (MV) sprayed 1 h prior to treatment. Photosystem II (PSII) activity (*Fv/Fm*) was measured using a Junior-PAM fluorometer (WALZ, Effeltrich, Germany).

### Recombinant enzymes

Recombinant proteins of AtGSTU1, AtGSTU2, AtGSTU3, AtGSTU4, AtGSTU5, AtGSTU7, AtGSTU8, AtGSTU10, AtGSTU13, AtGSTU17, AtGSTU18, AtGSTU19, AtGSTU22, AtGSTU23, AtGSTU24 and AtGSTU25, each fused to an *N*-terminal Strep-tag, were expressed in *Escherichia coli* and purified to homogeneity as described previously (Mano et al., 2017, 2019). These proteins are hereafter referred to as U1, U2, U3, U4, U5, U7, U8, U10, U13, U17, U18, U19, U22, U23, U24 and U25, respectively.

### Determination of peroxidase activity

Glutathione peroxidase (GPX) activity of GSTs was measured as described by Edwards & Dixon (2005), with modifications. The reaction mixture (1 mL) contained 550 μL of 0.25 M potassium phosphate buffer (pH 7.0), 100 μL glutathione reductase (6 U mL^−1^), 100 μL GSH (10 mM) and 100 μL NADPH (2.5 mM). After pre-incubation at 37 °C for 10 min, peroxide substrates were added: 100 μL of cumene-OOH (12 mM, aqueous) or 20 μL of HPODE mix (10 mM, ethanol; buffer volume adjusted accordingly). NADPH consumption was monitored at 340 nm for 1 min before and after adding 50 μL of GST solution. Enzyme activity was calculated from the change in NADPH consumption using the molar extinction coefficient 6220 M^−1^ cm^−1^.

### RCS treatment

RCS was administered to plants as vapor following Yamauchi et al. (2015). Arabidopsis plants (3 weeks old) were placed in a transparent plastic box (340 cm³; Nippon Genetics, Tokyo, Japan). A piece of paper towel containing an RCS solution in acetonitrile (3 μL total) was affixed to the lid, which was sealed immediately. This treatment corresponded to a vapor density of 10 nmol cm^−3^ upon complete evaporation. Plants were incubated at 25 °C for 30 min under light (80 µmol photons m^−2^ s^−1^). Control plants were exposed to acetonitrile vapor.

### Microarray analysis

Total RNA was extracted from at least six plants per treatment using an RNeasy Plant Mini Kit (Qiagen K.K., Tokyo, Japan). RNA quality was assessed by electrophoresis and spectrometry (A_260_/A_230_ and A_260_/A_280_ ratios > 1.8). Cy3-labelled cRNA was synthesized using the Low Input Quick Amp Labeling Kit (Agilent Technologies, Santa Clara, CA, USA). Microarray experiments were performed with an Agilent Arabidopsis v4.0 (44K) array using the one-colour method (Agilent Gene Expression Hybridization Kit and Wash Buffers Pack), according to the manufacturer’s protocols. Data were processed with Agilent Feature Extraction software, normalized, and clustered using the UPGMA method with Subio Platform software (Subio Inc., Kagoshima, Japan). Genes with expression ratios > 2.0 or < 0.5 were considered up- or downregulated, respectively. Reference expression datasets under various stress conditions were obtained from the AtGenExpress database (Kilian et al., 2007). Time-course data for shoots under heat, UV-B, drought and wounding (0.25–3 h) or oxidative, salt, osmotic and cold stress (0.5–12 h) were used to generate heat maps. Lists of up- and downregulated genes were compiled using Subio Platform software.

### Prediction of subcellular localization of GSTUs

Subcellular localization of each GSTU was predicted using SUBA5 platform (https://suba.live/) in which subcellular localization is determined by combination of data obtained from fluorescent protein tagging or mass spectrometry detection in subcellular purifications and by prediction using protein sequence features (Hooper et al. 2017; Hooper et al. 2022).

## Results

### Peroxidase activity of GST tau isoforms toward HPODE

When unsaturated fatty acids are oxidized by ROS, oxygen can be added at various carbon positions at the unsaturated bond(s), producing several positional hydroperoxide isomers. For example, oxidation of linoleic acid by hydroxyl radicals yields 9-HPODE and 13-HPODE, whereas oxidation by singlet oxygen generates not only these but also 10-HPODE and 12-HPODE (Triantaphylidès et al., 2008). Each positional isomer contains two C–C double bonds, and therefore exists as several geometric isomers. We prepared a mixture of HPODE positional isomers (hereafter referred to as ’HPODE mix’) by oxidizing linoleic acid with ROS and used it as the substrate to evaluate the LOOH-peroxidase activity of *A. thaliana* GSTUs. Among the 28 GSTUs in the genome, 16 isoforms that were successfully expressed in *E. coli* and purified were used in this study: U1, U2, U3, U4, U5, U7, U8, U10, U13, U17, U18, U19, U22, U23, U24 and U25 (Table 1). Peroxidase activity was quantified by monitoring the rate of glutathione oxidation, coupled to glutathione reductase activity, and expressed as the NADPH consumption rate.

**Table 1.** Comparison of GST tau isozymes from *A. thaliana* for their GSH-peroxidase activities. Isozyme proteins, expressed in *E. coli* and affinity-purified, were tested for peroxidase activity toward HPODE-mix and cumene-OOH as described in Materials and Methods.

| GST Tau# | Specific activity / nkatal (mg protein)<sup>-1</sup> |  | Activity ratio<br>(HPODE-mix/cumene-OOH) |
| --- | --- | --- | --- |
|  | HPODE-mix | cumene-OOH |  |
| 1 | 9.3 | 6.8 | 1.37 |
| 2 | 12.7 | 17.4 | 0.73 |
| 3 | 0.4 | 24.8 | 0.02 |
| 4 | 0.8 | 13.6 | 0.06 |
| 5 | 18.0 | 31.0 | 0.58 |
| 7 | 4.1 | 1.6 | 2.56 |
| 8 | 39.3 | 50.7 | 0.78 |
| 10 | 5.7 | 8.7 | 0.66 |
| 13 | 4.1 | 0.4 | 10.25 |
| 17 | 61.9 | 50.5 | 1.23 |
| 18 | 1.8 | 10.2 | 0.18 |
| 19 | 5.3 | 2.6 | 2.04 |
| 22 | 24.9 | 10.2 | 2.44 |
| 23 | 3.1 | 3.1 | 1.01 |
| 24 | 3.2 | 11.5 | 0.28 |
| 25 | 36.9 | 142.9 | 0.26 |

Specific activities toward cumene hydroperoxide (cumene-OOH) were highest for U25, U8 and U17, at 142.9, 50.7 and 50.2 nkatal mg^−1^, respectively. These were followed by U5 (31.0 nkatal mg^−1^), U3 (24.8 nkatal mg^−1^) and U2 (17.4 nkatal mg^−1^). By contrast, U13, U7, U19 and U23 displayed only marginal activity toward cumene-OOH. The distribution of isoform-specific activities was in good agreement with earlier findings (Dixon et al., 2009).

The highest activities toward HPODE mix were detected for U17, U8, U25 and U22, with specific activities of 61.9, 39.3, 36.9 and 24.9 nkatal mg^−1^, respectively, followed by U5, U2 and U1. By contrast, U3 and U4 showed substantially lower activity toward HPODE. Notably, the specific activity of U17 toward HPODE was comparable to the activity of GST theta 1–3 isoforms (97–170 nkatal mg^−1^) previously reported (Dixon et al., 2009).

Taken together, these results show that U2, U5, U8, U17, U22 and U25 accept both cumene-OOH and HPODE as substrates. U3 and U4 displayed a clear preference for cumene-OOH over HPODE, whereas U13 appeared to prefer HPODE over cumene-OOH (Table 1).

### AtGSTU17 utilizes multiple HPODE isomers as substrates

It remained unclear whether the peroxidase activity observed for each isoform against the HPODE mix reflected recognition of multiple molecular species simultaneously or selective recognition of particular isomers. To our knowledge, the degree to which plant GSTs discriminate among fatty acid hydroperoxide isomers has not previously been examined. We therefore isolated individual HPODE isomers—13EE-HPODE, 13ZE-HPODE, 12-HPODE, 10-HPODE and 9EE-HPODE—from the mixture (Ito et al., 2015b) and determined the enzymatic parameters of AtGSTU17, which showed the highest activity toward the HPODE mix.

U17 displayed Michaelis–Menten-type saturation curves for all five HPODE isomers at concentrations below 80 μM (Supporting Information Fig. S2). The Km values ranged from 8.97 μM (9EE-HPODE) to 24.65 μM (13EE-HPODE) (Table 2). To date, fatty acid hydroperoxides used for evaluating GPX or GST peroxidase activity have generally been tested as mixtures, without determination of kinetic parameters for individual isomers. Our analysis of purified HPODE isomers revealed, for the first time, that the Km values are approximately 10 μM.

**Table 2.** Kinetic properties of AtGSTU17 for various HPODE species.

| Substrate | Specific activity<br>(nmol s<sup>-1</sup> mg<sup>-1</sup>) | *k*<sub>cat</sub><br>(s<sup>-1</sup>) | *K*<sub>M</sub><br>(μM) | Catalytic efficiency<br>(s<sup>-1</sup> M<sup>-1</sup>) |
| --- | --- | --- | --- | --- |
| 9EE-HPODE | 40.37 | 1.13 | 8.97 | 126 × 10<sup>3</sup> |
| 10-HPODE | 52.16 | 1.46 | 9.27 | 158 × 10<sup>3</sup> |
| 12-HPODE | 96.81 | 2.71 | 20.4 | 133 × 10<sup>3</sup> |
| 13ZE-HPODE | 91.04 | 2.55 | 10.36 | 246 × 10<sup>3</sup> |
| 13EE-HPODE | 87.26 | 2.44 | 24.65 | 99.1 × 10<sup>3</sup> |

The k_cat_ values ranged from 1.13 s^−1^ (9EE-HPODE) to 2.71 s^−1^ (12-HPODE). The catalytic efficiencies (k_cat_/K_M_) varied from 99 s^−1^ mM^−1^ (3EE-HPODE) to 246 s^−1^ mM^−1^ (13ZE-HPODE), falling within a threefold range. These results demonstrate that U17 possesses broad substrate specificity toward various HPODE isomers.

### Substrate specificity profiling of AtGST tau isoforms

As shown above, AtGSTUs commonly exhibit peroxidase activity toward HPODE (Table 1). In addition, at least 10 of the 28 AtGSTUs have been shown to catalyse GSH conjugation with RCS, each isoform exhibiting distinct substrate specificities (Mano et al., 2019). For example, U19 and U24 display higher specific activities toward acrolein, whereas U17 and U18 preferentially conjugate HNE (Mano et al., 2017, 2019). By contrast, U10 and U25 do not show activity toward acrolein or HNE, but instead act on 2-hexenal or 12-oxo-phytodienoic acid (Dixon et al., 2009).

To comprehensively assess the substrate specificities of AtGSTUs, we generated substrate specificity profiles by integrating both peroxidase activity (substrates: HPODE, cumene-OOH) and RCS-conjugating activity (substrates: acrolein, crotonaldehyde, HNE and CDNB). For each substrate, the isoform with the highest specific activity among the 16 examined was assigned a relative activity value of 10. For example, for HPODE mix, GSTU17 was set at 10, GSTU25 exhibited a relative activity of 0.596, and GSTU3 had the lowest relative activity at 0.006. Isoforms that showed no detectable activity toward a substrate were assigned a value of 0.0001. These relative activities were plotted on radar charts in logarithmic scale to visualize isoform-specific substrate profiles (Fig. 2).

**Figure 1.**
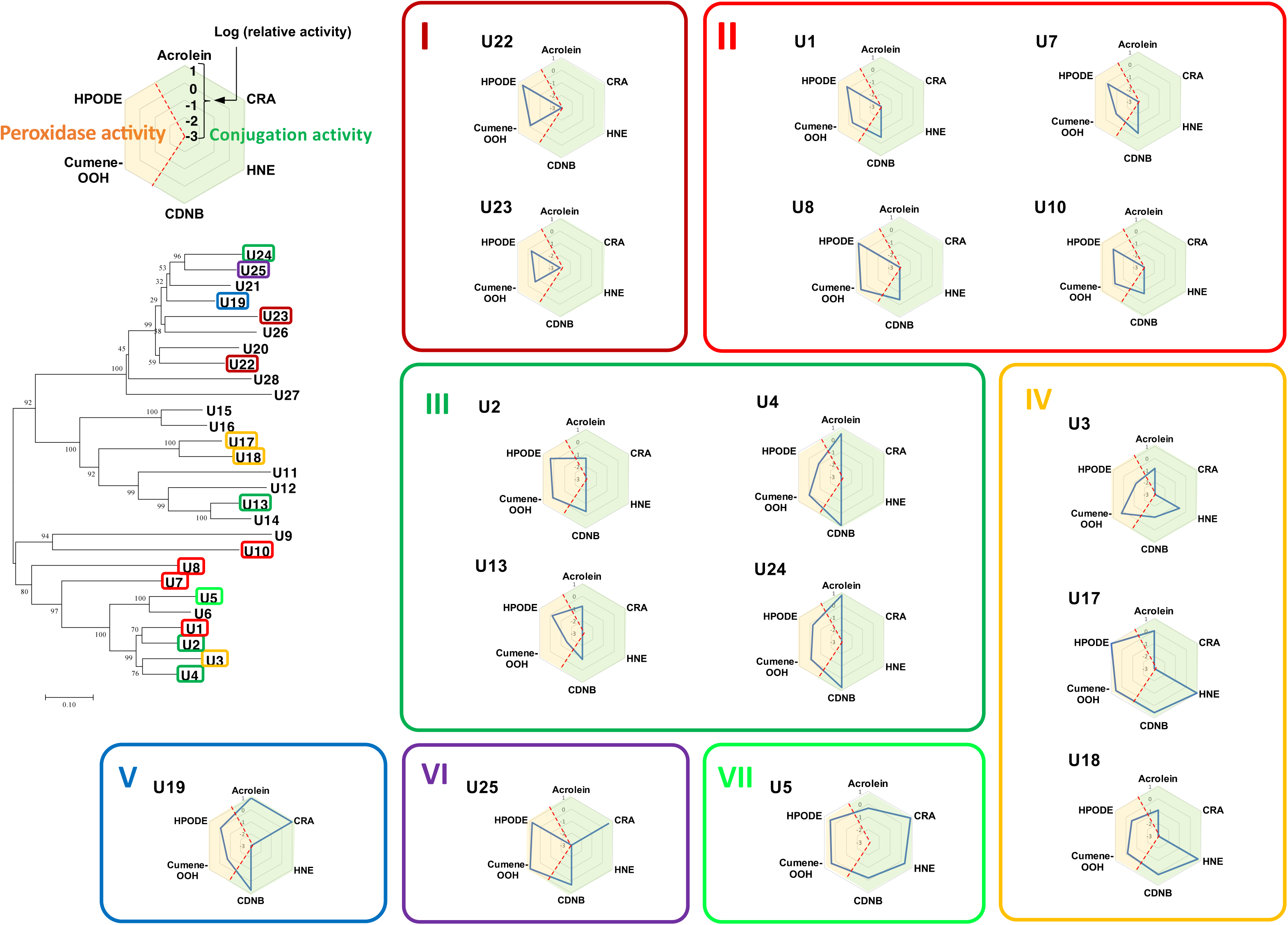
Substrate specificity profiles of AtGSTU isoforms. For each isoform, peroxidase activity toward cumene hydroperoxide (cumene-OOH) and HPODE mix (highlighted in light orange) and conjugation activity toward DCNB, acrolein, CRA and HNE (highlighted in light green) were normalized by setting the value of the most active isoform for each substrate to 10, and expressed as common logarithms in radar charts. Isoforms were classified into six substrate-specificity groups: Group I (dark red), peroxidase activity only; Group II (red), peroxidase activity plus conjugation activity only toward CDNB; Group III (green), peroxidase activity plus conjugation activity toward CDNB and acrolein; Group IV (orange), peroxidase activity plus conjugation activity toward acrolein and HNE; Group V (blue), peroxidase activity plus conjugation activity toward acrolein and CRA; Group VI (purple), peroxidase activity plus conjugation activity toward CRA. The phylogenetic tree (left) was constructed using amino acid sequences predicted from full-length cDNAs (obtained from https://www.arabidopsis.org/) with MEGA version 7.0.21 (https://megasoftware.net/).

**Figure 2.**
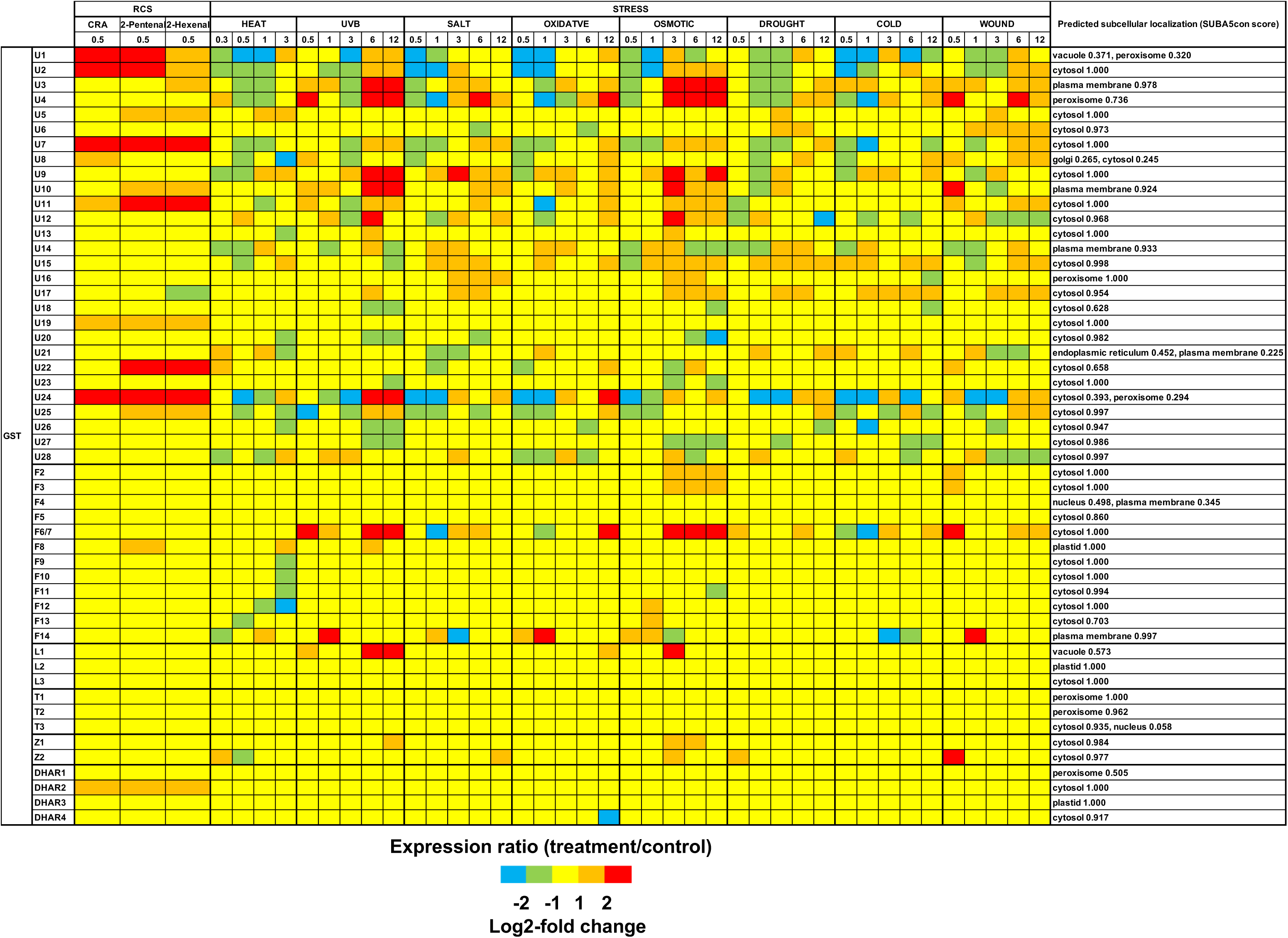
Genome-wide expression analysis of GST genes. Three-week-old *Arabidopsis thaliana* plants were exposed to volatile RCS (CRA, 2-pentenal, 2-hexenal) at a concentration of 10 nmol cm–3, and transcriptome analysis was performed focusing on GST classes. The fold induction relative to mock-treated controls (acetonitrile only) is shown as a heat map. Heat map data of stress responses were obtained from the public AtGenExpress dataset.

Based on these profiles, the 16 GSTUs could be classified into seven groups. Group I comprised isoforms (GSTU22, U23) exhibiting peroxidase activity toward HPODE but lacking conjugation activity. Group II contained isoforms with conjugation activity only toward the artificial substrate CDNB, with peroxidase activity predominating (GSTU1, U7, U8, U10). Group III included isoforms with HPODE peroxidase activity and conjugation activity toward acrolein (GSTU2, U4, U13, U24). Group IV comprised isoforms with HPODE peroxidase activity and conjugation activity toward acrolein and HNE (GSTU3, U17, U18). Group V contained GSTU19, which conjugated acrolein and crotonaldehyde, while Group VI contained GSTU25, which conjugated crotonaldehyde. Group VII comprised GSTU5, which exhibited conjugation activity toward acrolein, crotonaldehyde and HNE. Notably, this functional grouping did not strictly correspond to the phylogenetic clades of GSTUs (Fig. 2), suggesting that the functional plasticity of GST active sites enables divergent substrate specificities, as discussed below.

### Expression response analysis

Because reactive carbonyl species (RCS) are generated in planta under oxidative stress, they have been suggested to function as oxidative stress signals (Biswas & Mano, 2015; Biswas & Mano, 2016; Islam et al., 2016, 2020; Biswas et al., 2019). Representative examples include crotonaldehyde (CRA), produced non-enzymatically, and 2-hexenal, produced enzymatically (Yamauchi et al., 2015). To identify GSTU isoforms that are inducible by RCS, we analysed microarray datasets using CRA, 2-pentenal and 2-hexenal as stimuli.

Among the 28 GSTU isoforms, 12 were induced by RCS treatment (Fig. 2). Of these, U1, U2, U7, U11, U19 and U24 were induced by all three RCS tested. U5, U10, U22 and U25 were induced by 2-pentenal and 2-hexenal, U3 was induced only by 2-hexenal, and U9 was induced only by CRA. In contrast, other GST classes were not induced by RCS, with the exception of F8 and DHAR2.

Comparison with public stress-responsive expression data (AtGenExpress; Kilian et al., 2007) revealed that the RCS-inducible GSTUs (U1, U2, U3, U7, U10, U11, U24, U25) largely overlapped with isoforms induced by oxidative and osmotic stress. U5 and U22 resembled those induced by heat stress. By contrast, U19, which was induced by RCS, did not correspond to any known stress-responsive expression pattern.

### Subcellular localization of GSTU isoforms

Information on subcellular localization is essential for inferring the physiological functions of individual GSTU isoforms. We predicted the localization of each GSTU using the SUBA5 platform. As shown in the right panel of Fig. 2, the largest group, comprising 19 isoforms (U2, U5, U6, U7, U9, U11, U12, U13, U15, U17, U18, U19, U20, U22, U23, U25, U26, U27, U28), was predicted to localize to the cytoplasm. The plasma membrane was the second most common predicted location, with four isoforms (U3, U10, U14, U21), followed by the peroxisome with two isoforms (U4, U16). For U1, U8, U21 and U24, no reliable prediction of subcellular localization was obtained.

### Sensitivity of GSTU-deficient mutants to oxidative stress

The production of HPODE and the subsequent generation of RCS are promoted under oxidative stress conditions. If certain GSTU isoforms contribute to stress tolerance by detoxifying either RCS or HPODE, then their loss-of-function mutants would be expected to display enhanced sensitivity to oxidative stress. We therefore compared the oxidative stress sensitivity of GSTU-deficient T-DNA insertion mutants (*gstu2, gstu3, gstu4, gstu5, gstu6, gstu7, gstu17, gstu18, gstu19, gstu24, gstu25*) with that of wild-type (WT) plants. One of the major targets of RCS toxicity is photosystem II (PSII) (Yamauchi et al., 2011). Accordingly, we used the decline in PSII activity as a marker of oxidative stress sensitivity.

Under high-light (HL) stress, *gstu3, gstu17, gstu18* and *gstu25* exhibited significantly higher decreases in PSII activity compared with WT plants (Fig. 3b). When methyl viologen (MV) was applied as a stronger oxidative stressor, the differences in PSII damage between these mutants and WT were further amplified. In *gstu19*, PSII damage was not significantly different from WT under HL alone, but was higher than WT under HL plus MV treatment. By contrast, *gstu2, gstu4, gstu5, gstu6, gstu7* and *gstu24* did not show significant differences from Col-0 under either stress condition (Fig. 3c).

**Figure 3.**
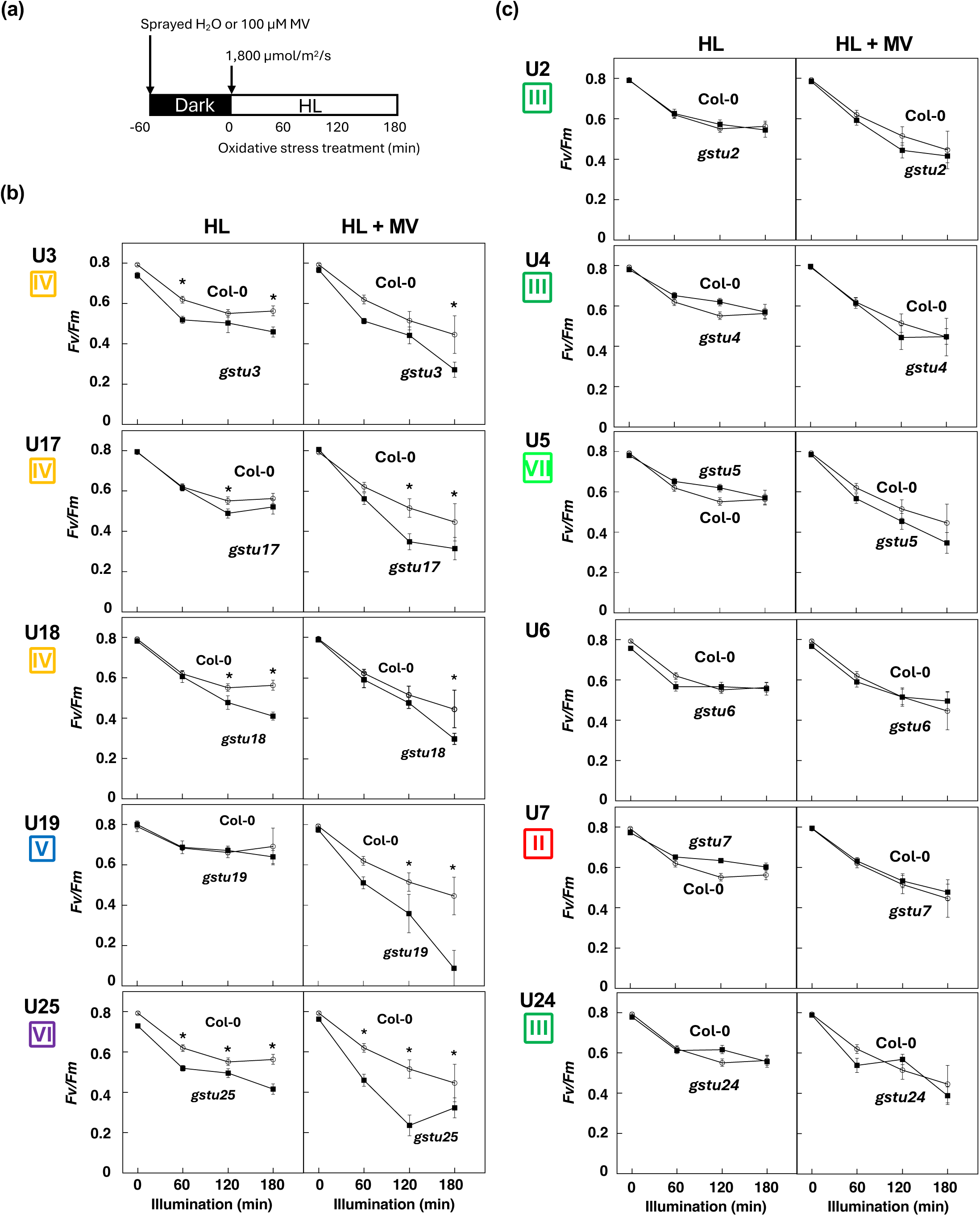
Effects of oxidative stress treatments on PSII activity in GSTU-deficient mutants. Three-week-old *Arabidopsis thaliana* plants were exposed to high light (HL) or high light plus methyl viologen (HL + MV), and PSII activity (Fv/Fm) was monitored over time. (a) Diagram of oxidative stress treatments. (b) GSTU mutants showing significantly greater PSII damage than wild type under oxidative stress. (c) Mutants showing no significant differences compared with Col-0 under the same treatments. *n* = 3; asterisks indicate significance at *P* < 0.05.

Among the GSTUs whose loss of function increased PSII oxidative damage, U3, U17 and U18 belong to Group IV (isoforms conjugating acrolein), U19 to Group V, and U25 to Group VI. By contrast, GSTUs from Groups II and III, which lacked RCS-conjugating activity, did not exhibit altered PSII sensitivity when disrupted. Likewise, disruption of GSTU5 (Group VII), which shows conjugating activity toward acrolein, CRA and HNE, did not affect oxidative stress sensitivity.

## Discussion

### Tau class GST isoforms exhibit peroxidase activity toward lipid peroxides

The *tau* class GST isoforms (GSTUs) are distributed only in land plants evolutionarily younger than tracheophytes, and within these species they represent the most diversified class of GST isoforms (Sylvestre-Gonon et al., 2019). This study demonstrates that *A. thaliana* GSTUs possess broad enzymatic capacity to counteract lipid oxidation stress through dual functions: glutathione-dependent reduction of LOOHs and conjugation of reactive carbonyl species (RCS). All 16 tested isoforms catalysed the GSH-dependent reduction of HPODE, indicating this activity is a conserved feature among GSTUs. As all of these isoforms had previously been reported to have peroxidase activity toward cumene-OOH (Dixon et al., 2009), this result was not unexpected. It should be noted, however, the relative ratios of activity toward cumene-OOH versus HPODE mix varied considerably among isoforms. For instance, U25 showed the highest activity toward cumene-OOH, whereas U17 displayed the highest activity toward HPODE mix (Table 1). Conversely, U3 and U4 exhibited very low activity toward HPODE mix but ranked sixth and seventh, respectively, in specific activity toward cumene-OOH among the 16 tested isoforms. These findings fill a long-standing gap in understanding how plant-specific GSTUs contribute to the detoxification of physiological LOOHs, which had previously been inferred mainly from assays using artificial peroxides such as cumene-OOH.

### Substrate selectivity and functional flexibility

Detailed kinetic analysis of GSTU17 revealed similar catalytic efficiencies for five distinct HPODE isomers, indicating broad substrate recognition of LOOHs. Such flexibility contrasts with the stringent substrate selectivity exhibited in its GSH-conjugation activity, where only HNE and acrolein serve, but crotonaldehyde and 2-hexenal not, as substrates (Mano et al., 2019). This combination of catalytic breadth and specificity implies a strategic bifunctionality: GSTU17, and likely other tau isoforms, may act as a general peroxidase removing diverse LOOHs generated non-enzymatically by reactive oxygen species (ROS), while simultaneously serving as a targeted detoxifier of highly cytotoxic RCS. The coexistence of these two activities within single enzymes provides an efficient mechanism to harness lipid peroxidation events at both the LOOH and carbonyl stages.

### Evolutionary conservation and diversification of catalytic properties

Phylogenetic and activity-based classification of 16 GSTU isoforms revealed that functional groupings defined by substrate preferences did not coincide with sequence-based clades (Fig. 2). This discrepancy underscores the catalytic plasticity of GST active sites, consistent with previous reports that minor amino acid substitutions can drastically shift substrate specificity or catalytic mode (Dixon, Davis & Edwards, 2002; Axarli et al., 2024). During land plant evolution, the peroxidase function of GSTUs may have been conserved as an ancestral enzymatic trait, while conjugation activity toward particular RCS evolved independently among lineages. Such diversification would have been favoured under environmental conditions that imposed oxidative challenges on early terrestrial plants. Isoforms combining high peroxidase activity with RCS-conjugation potential would provide selective advantages by mitigating lipid oxidation-derived cytotoxicity. The present data therefore support the view that the *tau* class evolved as a versatile detoxification system that integrates multiple layers of lipid peroxide metabolism.

### RCS as inducers of *GSTU* expression

RCS, including crotonaldehyde, 2-pentenal and 2-hexenal, not only act as toxic by-products of lipid peroxidation but also serve as electrophilic signals activating antioxidant defence genes (Yamauchi et al., 2015). Transcriptome analysis identified twelve GSTU genes strongly induced by these RCS, whereas other GST classes (*phi*, *lambda*, *theta*, *zeta* and DHAR) showed little or no response (Fig. 2). This selective responsiveness implies that GSTUs form a core component of an RCS-responsive regulatory circuit. Such RCS-mediated feed-forward regulation is likely to represent an adaptive response that strengthens protection against lipid oxidation stress.

Interestingly, the expression patterns of RCS-inducible GSTUs (U1, U2, U3, U5, U7, U10, U11, U22, U24, U25) overlapped extensively with those induced by oxidative, osmotic and heat stress, supporting the notion that RCS function as endogenous mediators linking ROS generation to transcriptional activation of stress-protective genes (Fig. 2). Among these, U17 was uniquely downregulated by 2-hexenal, consistent with earlier findings that its overexpression compromises drought and salt tolerance by depleting the glutathione (GSH) pool (Chen et al., 2012). This inverse regulation may represent an adaptive mechanism that maintains GSH homeostasis under sustained stress.

Many proteins critical for cellular homeostasis form multigene families consisting of both constitutively expressed members and inducible members that function under stress. Examples include heat shock proteins (HSPs) involved in heat responses (Kozeko, 2021) and superoxide dismutases (SODs) involved in antioxidant defense (Huo et al., 2022). Similarly, GSTs can be broadly divided into constitutively expressed groups (*phi*, *lambda*, *theta*, DHAR) and stress-inducible groups, particularly the GSTUs, which themselves display diverse expression patterns. This indicates that GSTUs, like HSPs and SODs, represent a crucial enzyme family for maintaining cellular homeostasis.

### Physiological relevance of RCS detoxification

Loss-of-function analysis provided direct evidence that specific GSTUs contribute to oxidative stress tolerance. Mutants deficient in GSTU3, GSTU17, GSTU18, GSTU19 or GSTU25—isoforms with pronounced RCS-conjugating activity—exhibited enhanced photoinhibition damage under high-light or MV treatment (Fig. 3). In contrast, mutants of GSTUs possessing only peroxidase activity (such as U2 or U7) showed no change in stress sensitivity. These results indicate that RCS detoxification, rather than LOOH reduction per se, is the dominant determinant of the tolerance against photooxidative stress.

Predicted subcellular localisations further refine this interpretation. U3 localizes to the plasma membrane, while U17, U18, U19 and U25 are cytosolic (Fig. 2). The enhanced stress sensitivity observed in the respective mutants suggests that cytoplasmic RCS detoxification is critical for oxidative stress tolerance. During MV treatment, chloroplasts act as the major ROS source, producing H₂O₂ that diffuse into the cytoplasm where it initiates secondary lipid peroxidation in plasma membrane. The enhanced stress sensitivity of cytoplasmic GSTU mutants suggests that the detoxification of RCS in the cytosol and at the plasma membrane is essential as a front-line defence for buffering cytoplasmic and membrane environments against lipid oxidation stress originating from chloroplast ROS.

Collectively, our findings reveal that plant *tau* class GSTs have evolved as pivotal enzymes bridging ROS metabolism, lipid peroxide reduction, and RCS detoxification. Their dual catalytic capacity enables plants to intercept the lipid oxidation cascade at multiple stages, while RCS-responsive transcription ensures dynamic up-regulation of protective isoforms under stress. This integrated network provides a molecular basis for the remarkable oxidative resilience of terrestrial plants. Further characterization of both RCS- and lipid peroxide-detoxifying activities will be essential for elucidating the physiological functions of this enzyme family.

## Acknowledgements

This work was supported by JSPS KAKENHI (Grant Numbers 20H03278, 24H00504, 15612760 and 22485763), by the YU Project for Formation of the Core Research Center from Yamaguchi University, and by Cross-cutting Research Projects from the United Graduate School of Agricultural Science, Tottori University.

## Author contributions

YY and JM designed research; NM, FK, CS, RN and JI performed research; KN, MM, YS analyzed data; YY, JI and JM wrote the paper.

## Data availability

All the data and materials integral to this study are available within this article and Supporting Information. The microarray data have been deposited in the Gene Expression Omnibus under the accession number GEO: GSM1569068.

