## Supporting Information Fig. S1 for "Plant-specific *tau* class glutathione transferases safeguard against lipid oxidation stress through dual detoxification of lipid peroxides and reactive carbonyl species"

9-10*E*,12*E*-HPODE
(9EE-HPODE)

9-10*E*,12*Z*-HPODE
(9EZ-HPODE)

10-8*E*,12*Z*-HPODE
(10-HPODE)

12-9Z,13*E*-HPODE
(12-HPODE)

13-9*E*,11*E*-HPODE
(13EE-HPODE)

13-9Z,11*E*-HPODE
(13ZE-HPODE)

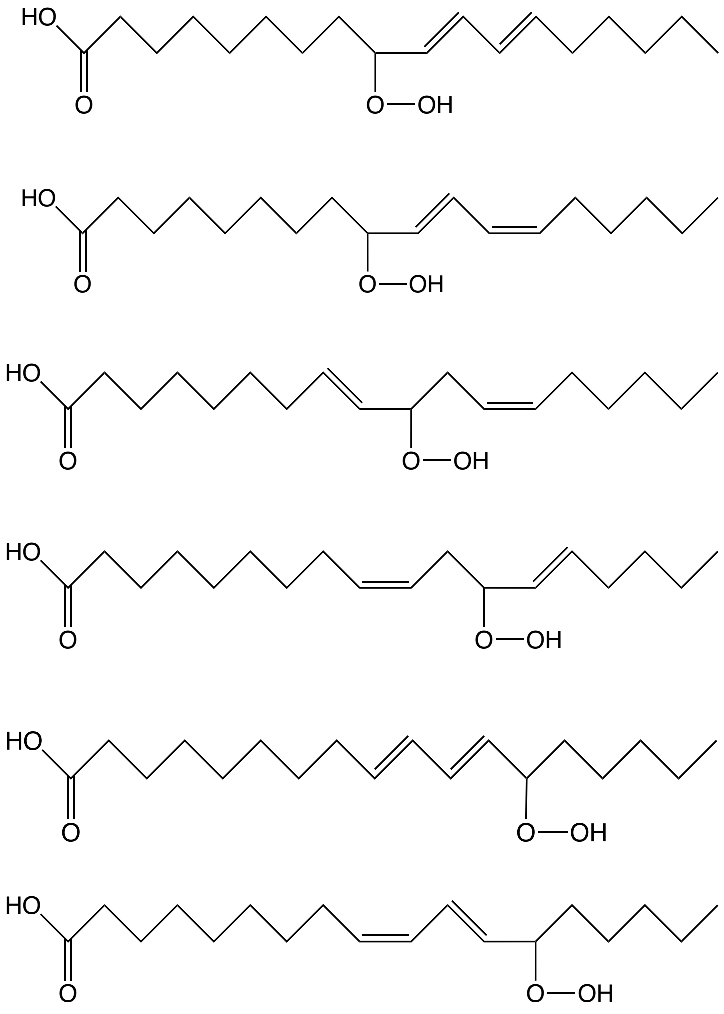

Supplementary Figure S1. Structure of HPODE isomers.
