## Supporting Information Fig. S2 for "Plant-specific *tau* class glutathione transferases safeguard against lipid oxidation stress through dual detoxification of lipid peroxides and reactive carbonyl species"

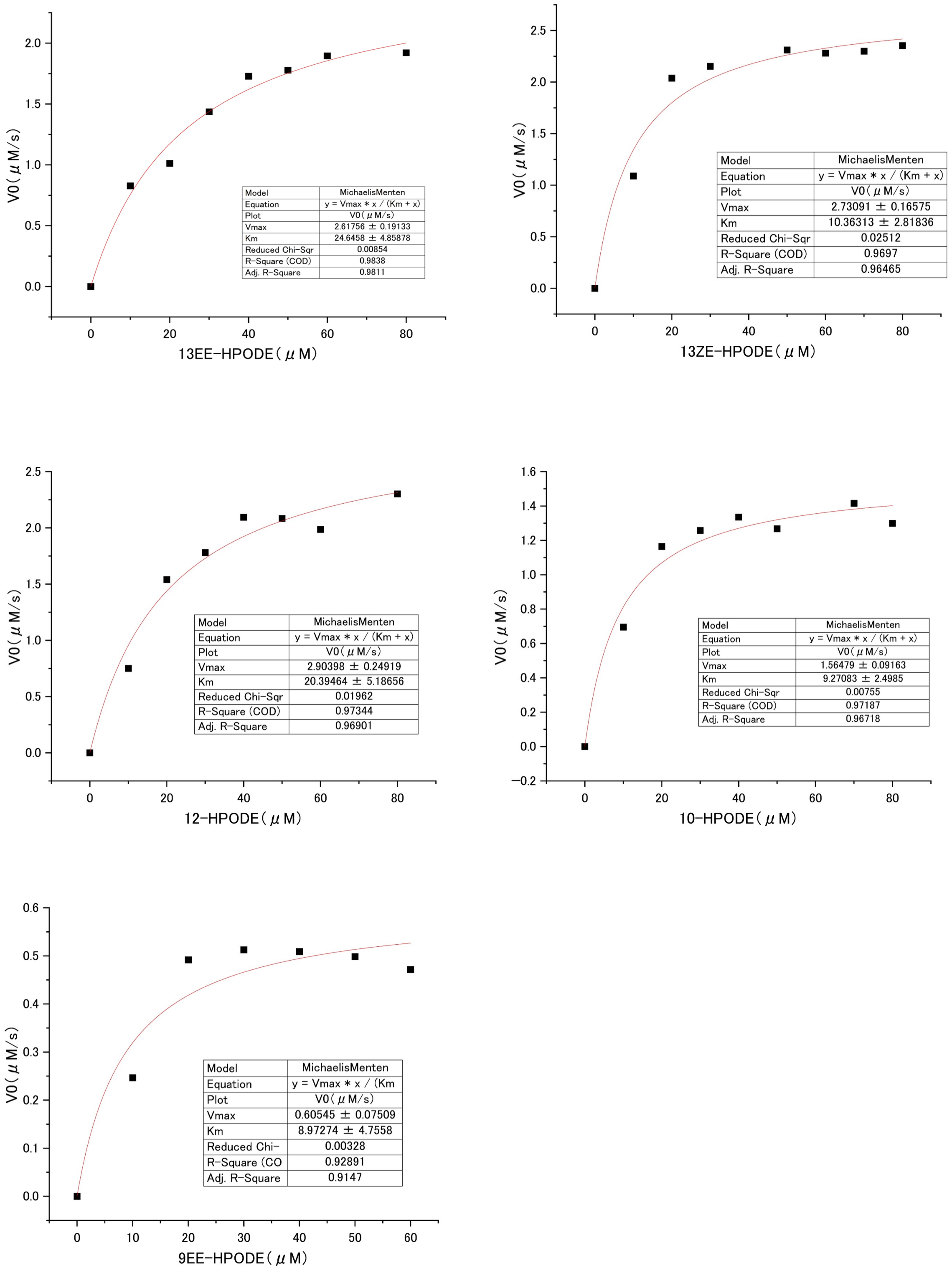


Supplementary Figure S2. Kinetic properties of AtGSTU17 for 13EE-HOPDE, 13ZE-HPODE, 12-HPODE, 10-HPODE and 9EE-HPODE. Reaction rate was determined as in Materials and Methods. Purified HPODEs were dissolved in ethanol and used as substrates. Final ethanol concentration in the reaction mixture was 2% (v/v). Each data point was an average of two assays.
