## Supporting Information Table S1 for "Plant-specific *tau* class glutathione transferases safeguard against lipid oxidation stress through dual detoxification of lipid peroxides and reactive carbonyl species"

Supplementary Table S1. Primers used for Homo-Hetero selection of T-DNA tagged mutants.

| Primer | Sequence | SALK ID |
|---|---|---|
| GSTU2 LP | TGCACCCATTAATTGATTTCC | 062920C |
| GSTU2 RP | ACCAAAGTGACCTCAGCAGTG | 062920C |
| GSTU3 LP | TATCCAATGAACCGTGAACAC | 17999 |
| GSTU3 RP | GTCCATATGACAAAGCCATGG | 17999 |
| GSTU4 LP | TAATTCACTTTTGGACCACGC | 143928C |
| GSTU4 RP | AGTCATCGGTGGTTCTAGGTG | 143928C |
| GSTU5 LP | ATCGTGAGGTGGTGAGTCTTG | 107148 |
| GSTU5 RP | TTCCTTTCTCGTCTGCTCTTG | 107148 |
| GSTU6 LP | CTATTGCCGGATTAGCTAGGG | 207079C |
| GSTU6 RP | TGGCTCTCAAACTCAAAGGTG | 207079C |
| GSTU7 LP | GAAACAATCTCTTCGTGATTTCC | 086642C |
| GSTU7 RP | CAAATCTCTCGTCGCTTCAAC | 086642C |
| GSTU17 LP | GCCACGCATATAAAGAAATGC | 139615C |
| GSTU17 RP | GCTGACTGCTAACTCGGTGAC | 139615C |
| GSTU18 LP | TTGGCCACACTTCAATTTCTC | 096297C |
| GSTU18 RP | TTTCCTTGGCTACAATCGTTG | 096297C |
| GSTU19 LP | TACATAGCCAAAGTCATCGCC | 041942 |
| GSTU19 RP | TTTAGCGATCGTAACAATGGC | 041942 |
| GSTU24 LP | CAGTGGCCTAAGCAACTAACG | 034472C |
| GSTU24 RP | CTGCAAACCAGCTATGGAATC | 034472C |
| GSTU25 LP | ATTTGGCACTCCACCAAAATC | 042213 |
| GSTU25 RP | AATCAAGAACAGCAATGGCAG | 042213 |
| pROK2 LBb1.3* | ATTTTGCCGATTTCGGAAC | |
| pROK2 LBa1* | TGGTTCACGTAGTGGGCCATCG | |

*Left border primers of the T-DNA insertion used for Homo line selection
