## Supporting Information Table S2 for "Plant-specific *tau* class glutathione transferases safeguard against lipid oxidation stress through dual detoxification of lipid peroxides and reactive carbonyl species"

Supplementary Table S2. Specific activities of GSTUs.

| | Specific activity (nkat/mg) | | | | | |
|---|---|---|---|---|---|---|
| | Acrolein | CRA | HNE | CDNB | Cumene-OOH | HPODE mix |
| U1 | < 0.01 | - | < 0.01 | 8.83 | 6.80 | 9.30 |
| U2 | 0.24 | < 0.01 | < 0.01 | 56.0 | 17.4 | 12.7 |
| U3 | 0.72 | < 0.01 | 0.94 | 2.40 | 24.8 | 0.40 |
| U4 | 15.0 | < 0.01 | < 0.01 | 275 | 13.6 | 0.80 |
| U5 | 2.38 | 0.83 | 8.17 | 44.5 | 31.0 | 18.0 |
| U6 | < 0.01 | < 0.01 | < 0.01 | 9.48 | - | - |
| U7 | < 0.01 | - | < 0.01 | 10.8 | 1.60 | 4.10 |
| U8 | < 0.01 | < 0.01 | < 0.01 | 12.2 | 50.7 | 39.3 |
| U9 | - | - | - | - | - | - |
| U10 | < 0.01 | < 0.01 | < 0.01 | 4.55 | 8.70 | 5.70 |
| U11 | < 0.01 | < 0.01 | - | < 0.01 | - | - |
| U12 | - | - | - | - | - | - |
| U13 | 0.81 | < 0.01 | - | 3.83 | 0.40 | 4.10 |
| U14 | - | < 0.01 | - | 0.24 | - | - |
| U15 | - | - | - | - | - | - |
| U16 | < 0.01 | < 0.01 | < 0.01 | 84.0 | - | - |
| U17 | 5.17 | < 0.01 | 39.0 | 102 | 50.5 | 61.9 |
| U18 | 0.62 | < 0.01 | 21.2 | 36.5 | 10.2 | 1.80 |
| U19 | 50.7 | 1.20 | < 0.01 | 152 | 2.60 | 5.30 |
| U20 | < 0.01 | < 0.01 | < 0.01 | 30.2 | - | - |
| U21 | - | - | - | - | - | - |
| U22 | - | - | - | - | 10.2 | 24.9 |
| U23 | < 0.01 | < 0.01 | < 0.01 | < 0.01 | 3.10 | 3.10 |
| U24 | 34.2 | < 0.01 | < 0.01 | 158 | 11.5 | 3.20 |
| U25 | < 0.01 | 0.47 | < 0.01 | 58.7 | 143 | 36.9 |
| U26 | - | < 0.01 | 9.50 | 1.24 | - | - |
| U27 | < 0.01 | < 0.01 | < 0.01 | 414 | - | - |
| U28 | - | < 0.01 | 1.00 | 286 | - | - |
